# Sperm length ornamentation associates with survival and sterility after heatwaves among *Drosophila* species

**DOI:** 10.64898/2026.10.08.757264

**Authors:** Ana Marquez-Rosado, Liam R. Dougherty, Won-Yong Choi, R. Antonio Gomez, Christopher Mitchell, Scott Pitnick, Stuart Wigby

## Abstract

Whether sexual selection facilitates or hinders adaptation to novel environments is a question of fundamental importance due to current rapid environmental change. Using phylogenetic comparative analysis in 42 *Drosophila* species, we found that sperm length – a sexually selected ornament – positively associates with the ability of males to survive high temperatures. However, the ability to remain fertile does not increase with sperm length to the same extent, meaning that longer sperm species experience a larger gap between temperatures that kill and temperatures that sterilize males. The findings are consistent with the idea that strong sexual selection on condition-dependent exaggerated traits may facilitate somatic resilience to environmental stressors, but that susceptibility to heat-induced infertility may be a cost of extreme sperm ornamentation. Overall, the data suggests multifaceted links between postcopulatory sexual selection and climate change.

## Introduction

Extreme heat events are increasing in frequency and severity, posing a severe threat to biodiversity as temperatures exceed survivable limits. A key current challenge is to understand the capacity of organisms to adapt to novel thermal environments. Sexual selection could play a role in enhancing or constraining adaptation to changing environments[1]. For example, the growth and maintenance of male sexual ornaments is likely to be condition-dependent and thus linked to genes that underpin core survival traits[2, 3]. However, because extreme ornaments should be costly and consume resources, they are commonly thought to trade-off against other fitness traits, which could include those key to environmental adaptation[4, 5]. Therefore, the extent to which sexual selection facilitates or hinders adaptation to environmental stress remains contentious[1].

To explore this problem, we examined the relationship between a sexually selected ornament and resilience to heat stress across 42 *Drosophila* species. Recent work in this genus has demonstrated that sexual selection favors extreme sperm length. As a target of postcopulatory sexual selection, sperm length has evolved dramatically among *Drosophila* species and is now recognized as one of nature’s most extreme ornaments[4]. There is strong evidence that sperm length is condition-dependent: only high-quality males can produce abundant sperm, and this effect is more pronounced in species with longer sperm[4]. In *D. melanogaster*, genetic variation in sperm length is highly pleiotropic – sperm length genes contribute to metabolic function, nervous system development, and sensory biology[6]. As such, sperm length has the potential to reflect male genetic quality[6], which could relate to the ability to withstand stressful conditions[2]. Honest signals should be costly; consistent with this prediction, *Tribolium* produces shorter sperm under larval nutritional stress[7], and *Drosophila* species with longer sperm have delayed sexual maturity and reduced sperm numbers[4, 5].

Animal sperm are especially sensitive to heat, such that the males of many species often become infertile at temperatures lower than the Lethal Temperature[8]. This is true for many drosophilids[8] and other invertebrates[9] but also mammals, including humans, where recent evidence shows that longer heat wave exposure reduces semen quality and lowers sperm count[10]. The present study was fueled by Parratt et al.’s finding that the temperature that causes infertility (‘Thermal Fertility Limit’) better predicts species distribution with increasing temperatures than the temperature that causes death (‘Lethal Temperature’)[8]. We examined the among-species relationships between sperm length, Lethal Temperature, and Thermal Fertility Limit[8] via phylogenetic generalized least squares analyses to disentangle whether longer-sperm species are more vulnerable to heat.

## Materials and Methods

We conducted phylogenetic generalized least squares analysis using Thermal Fertility Limits and Lethal Temperatures reported for 41 species by Parratt et al. 2021, plus our own new data for *D. pachea[8]*. We obtained sperm length data from the SpermTree Database and Syed et al. 2025[6, 18]. For those species with polymorphic sperm we used the fertilizing morph (which is always longer). All analyses were performed using R version 4.2.2[19]. We used a pruned *Drosophila* phylogeny published by Kim et al.2024[20]. Sperm length was log-transformed, and for a given species the Thermal Fertility Limit was constrained to not be higher than the Lethal Temperature. Multicollinearity between variables was checked by running a principal component analysis corrected for phylogeny using phyl.pca() from *phytools* 2.4-4[21].

We first calculated the phylogenetic signal for each trait using the maximum-likelihood approach (Pagel’s λ) with the phylosig() function from the *phytools* package[21]. Then, three models were built to test how sperm relates to either Lethal Temperature, Thermal Fertility Limit, or the Lethal Temperature – Thermal Fertility Limit Gap using phylogenetic generalized least squares regression. We ran the regressions using the phylolm() function from the *phylolm* package in R[22]. For each model, we transformed the variance-covariance matrix using the parameters lambda, kappa, delta, or alpha to fit the data by maximum likelihood under different evolutionary models[23, 24]. These parameters were directly implemented in the phylogenetic generalized least squares calculations of phylogenetic signal in the residuals, and each of them represents a different generating model for the error in distances among species[25]. By comparing them, we more accurately capture the degree of phylogenetic dependence in the data and rigorously evaluate basic models of the underlying evolutionary process. Models were compared using AICc scores. Assumptions of normality and homoscedasticity of the residuals were verified in all models.

We next used phylogenetically-corrected path analysis to search for causal evolutionary links between sperm length, Lethal Temperature, Thermal Fertility Limits (Thermal Fertility Limit), and the Lethal Temperature – Thermal Fertility Limit Gap[26]. We used the *phylopath* package version 1.3.1 in R[27], which is based on the d-separation method by Shipley[26] and controls for phylogeny as explained by von Hardenberg and Gonzalez-Voyer[28]. Using previous knowledge, we established 8 a priori hypotheses (**Fig. 1**, paths I-VIII) based on theoretical hypotheses that link somatic and gametic mechanisms of thermal tolerance, sexual selection and climate. The hypotheses are as follows: (I). **Sperm Length → Thermal Fertility Limit**: Sperm length and Thermal fertility limit are negatively related because long sperm are more sensitive to heat due to an increased duration of spermatogenesis and delayed sexual maturity, increasing the window for heat-related damage; (II). **Sperm Length → Thermal Fertility Limit – Lethal Temperature Gap**: Sperm length and the Lethal Temperature – Thermal Fertility Limit Gap are positively related because long sperm are more sensitive to heat, increasing the probability that heat-related damage in spermatogenesis will occur at a lower temperature than the animal can survive; (III). **Thermal Fertility Limit → Lethal Temperature**: There is a positive relationship between these traits because the genetic mechanisms of heat resistance for long sperm are shared with the soma; (IV). **Lethal Temperature → Thermal Fertility Limit**: Lethal temperature and Thermal Fertility limit are positively related because the genetic mechanisms of heat resistance for the soma are shared with long sperm; (V). **Thermal Fertility Limit → Thermal Fertility Limit – Lethal Temperature Gap**: These traits are negatively related because species adapted to hotter environments have better sperm heat-resistance mechanisms and have a smaller window for heat damage during spermatogenesis; (VI). **Lethal Temperature → Thermal Fertility Limit – Lethal Temperature Gap**: There is a positive relationship because sperm heat tolerance is constrained and lags behind somatic heat tolerance; (VII). **Lethal Temperature → Sperm Length**: Lethal temperature and sperm length are positively correlated because in hotter environments, increases post-copulatory sexual selection, which increases females’ remating rate, leading to sexual selection for longer sperm; (VIII). **Sperm Length** → **Lethal Temperature:** There is a positive relationship because high post-copulatory sexual selection, which drives longer sperm via processes that favour better overall genetic quality, will boost stress resistance. Based on these hypotheses, we constructed 9 models of causal relationships among the variables of interest. Then, models were compared using CIC (C-statistic information criterion) which is analogous to Akaike Information Criterion and estimates the goodness of fit of the models. Any models with a ΔCIC of 2 or less are statistically indistinguishable and are equally likely to be the best-fitting model. Of the 9 models, 4 fitted the data equally well, we therefore averaged the coefficients from all best-fitting models to get the final predicted values.

**Figure 1.**
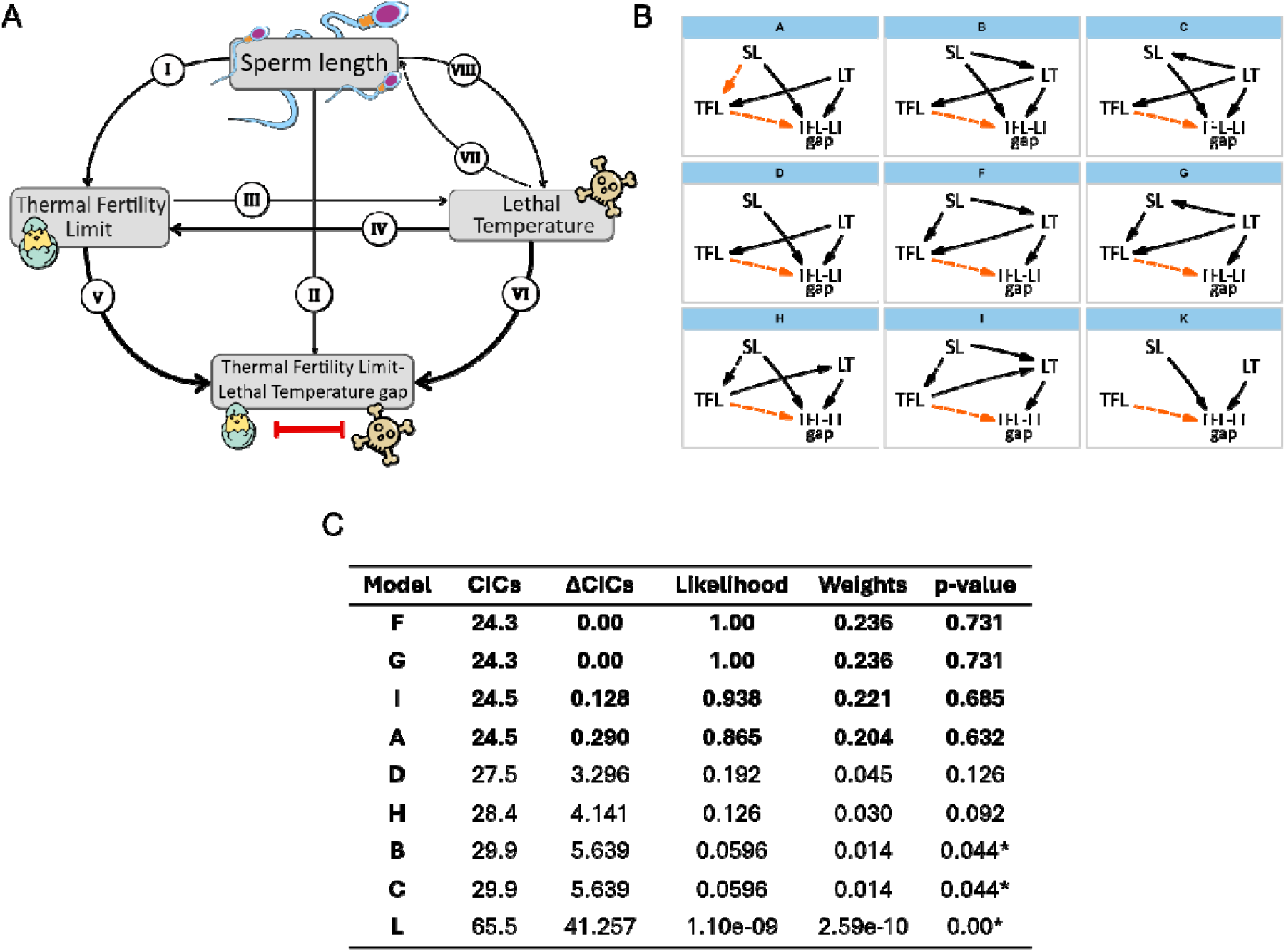
Path analysis methods. A) Path analysis full model showing all potential causal links between sperm length. Thermal Fertility Limit, Lethal Temperature, and the Thermal Fertility Limit – Lethal Temperature gap, based on theories linking gametic and somatic mechanisms of thermal tolerance and sexual selection. The direction of arrows indicates causality. B) Nine causal models were established based on the hypothesis between the variables of interest. The arrow indicates causality, and the orange-dashed lines represent a negative association between traits. SL: Sperm Length, LT: Lethal Temperature, TFL: Thermal Fertility Limit and TFL-LT gap: Thermal Fertility Limit – Lethal Temperature gap. C) Path Analysis results. Models are ranked according to their ΔCICs. When the p-value is significant, it means a rejection of that model. Models in bold letters indicate ΔCICs ≤ 2.

## Results

We found that sperm length was significantly positively associated with Lethal Temperature (t=2.713, p=0.010, F=7.359, r^2^=0.155, α=0.070, coefficient=0.627±0.231; α being the phylogenetic signal of the residuals) but not with Thermal Fertility Limit (t=1.239, p=0.222, F=1.536, r^2^=0.037, α=0.151, coefficient=0.344±0.278; α being the phylogenetic signal of the residuals; **Fig. 2a and c**). There was also a significant positive relationship between sperm length and the difference between Thermal Fertility Limit and Lethal Temperature (Thermal Fertility Limit – Lethal Temperature ‘Gap’[8]; t=2.071, p=0.045, F=4.288, r2=0.097, λ=0.349, coefficient=0.425±0.205; being λ the phylogenetic signal of the residuals) across species (**Fig. 2b**). Hence, as sperm length increases across species, so does the ability, on average, to survive episodes of extreme heat. However, because the Thermal Fertility Limit does not concomitantly increase, the probability that sub-lethal heat can render males infertile also increases. The trait phylogenetic signal for sperm length is λ=0.928 (p<0.01), λ=0.460 (p<0.01) for Lethal Temperature, λ=0.514 (p<0.01) for Lethal Temperature-Thermal Fertility Limit Gap and λ=887 (p=0.1) for Thermal Fertility Limit.

**Figure 2.**
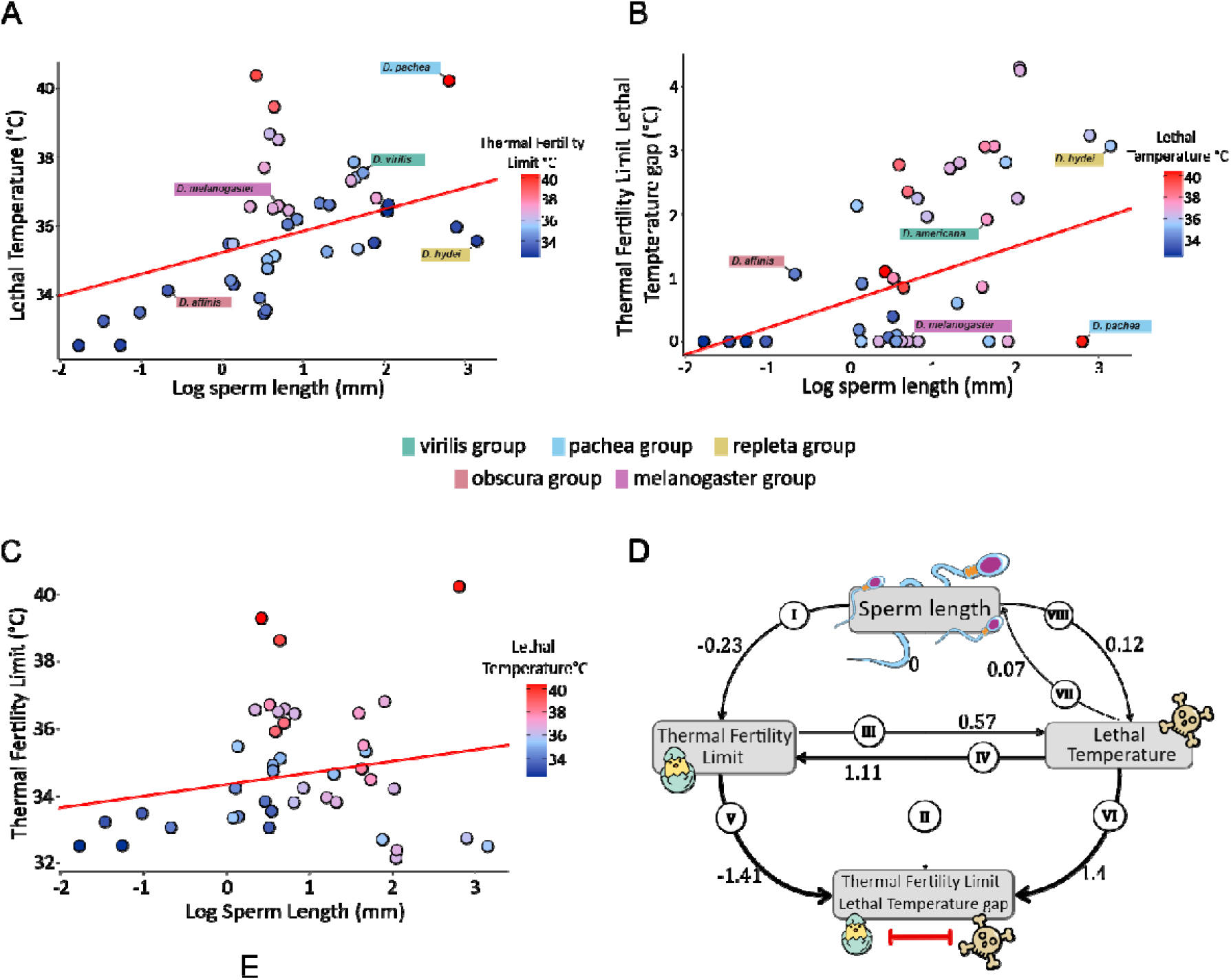
Association of sperm length and thermal traits. A) Linear correlation of log sperm length and Lethal Temperature under Ornstein-Uhlenbeck model. B) Linear correlation of log sperm length and Lethal Temperature-Thermal Fertility Limit Gap under Pagel’s lambda model. C) Linear correlation of log sperm length and Thermal Fertility Limit under Ornstein-Uhlenbeck model. For A, B, and C, each dot represents a species and is colour-coded by the species’ Thermal Fertility Limit or Lethal Temperature, a measure of heat tolerance. D) Standardized path coefficients of the averaged model. The direction of the arrows indicates evolutionary causality, and the thickness of each arrow represents the strength of the effect (larger positive or negative values indicate stronger effects).

To examine the underlying causal structure relating sperm length to Lethal Temperature, Thermal Fertility Limit, and the Thermal Fertility Limit – Lethal Temperature Gap, we used a phylogenetically corrected path analysis approach. Consistent with the phylogenetic generalized least squares analysis (**Fig. 2a**), the resulting model (**Fig. 2d**) indicates the Lethal Temperature is strongly influenced by sperm length (path VIII) and, to a lesser extent, these two variables are associated in the opposite direction (path VII). Sperm length is also negatively associated with the Thermal Fertility Limit (path I), while the Lethal Temperature – Thermal Fertility Limit Gap is related to sperm length indirectly, via opposing effects on the Lethal Temperature and Thermal Fertility Limit. As expected, the Lethal Temperature and Thermal Fertility Limit have opposing signs (paths V and VI) because the Thermal Fertility Limit can never be higher than the Lethal Temperature, and a wider gap can only arise from a lower Thermal Fertility Limit (negative coefficient) and/or a higher Lethal Temperature (positive coefficient).

## Discussion

Our results suggest that sperm length relates to among-species variation in the upper limits of thermal stress tolerance. Path VIII (**Fig. 2d**) – which suggests that sperm length drives increases in Lethal Temperature – is based on the hypothesis that post-copulatory sexual selection has favored the evolution of longer sperm because they confer indirect genetic benefits related to the ability to maintain essential biological processes when facing environmental challenges[2, 6]. A potential mechanism underpinning this could be shared developmental pathways between sperm and somatic organs that underpin heat survival. In *D. melanogaster*, sperm length genes show a bias towards developmental and central nervous system expression[6]. In turn, the central nervous system is known to be important in heat responses: heat causes spreading depolarisation in the adult central nervous system associated with the loss of coordination[11], while in larvae, heat-induced death is genetically regulated by the *Phae1* gene, which is highly expressed in the central nervous system [12]. In further support of the contention that sperm length reflects genetic condition, it is significantly positively correlated with starvation resistance and lifetime female fecundity across isogenic lines of *D. melanogaster[6*]. It is important to note, however, that no equivalent studies have been conducted with other *Drosophila* species.

Path analyses also suggest that the Lethal Temperature may drive sperm length, albeit to a lesser extent than *vice versa* (**Fig. 2d, path VII**). Some studies suggest that *D. melanogaster* females increase polyandry in hotter environments due to positive effects on fecundity and offspring[13] which could in turn increase the intensity of post-copulatory sexual selection for longer sperm. Note, however, that evidence of increasing polyandry with temperature is not universal[14].

The negative association between sperm length and Thermal Fertility Limit, and the lack of direct relationship between sperm length and Lethal Temperature – Thermal Fertility Limit Gap indicated by the path analysis (**Fig. 2d, path I**), appear to contrast with the phylogenetic generalized least squares results (**Fig 2b-c**). However, taken as a whole, the analyses underscore the fact that the association between sperm length and the Lethal Temperature – Thermal Fertility Limit Gap revealed by phylogenetic generalized least squares derives from the different relationships between sperm length and Lethal Temperature *versus* sperm length and Thermal Fertility Limit. We hypothesize that an increased duration of spermatogenesis[15, 16] and time to male reproductive maturity[5] in longer sperm species may create a greater window of opportunity for heat-related damage during critical stages of sperm production[17], increasing the probability of a Lethal Temperature – Thermal Fertility Limit Gap.

Our results are consistent with the theory positing that sexually selected ornaments (e.g., sperm length in *Drosophila*;[4, 6]) are linked to survival under environmental stress. The most parsimonious mechanism, based on current knowledge, is condition dependence of sperm length, and selective processes that pleiotropically link ornament genes with those underlying core somatic development and metabolic functions[2, 3]. We note, however, that any sexual selection processes that favor longer sperm that also improve population fitness via stress resistance traits[1] could generate the patterns seen in our data. There is no evidence to suggest that sperm length, *per se*, relates to thermal tolerance, nor that females have evolved fertilization bias in favor of longer sperm because it conveys thermal tolerance. Rather, our results support the contention that *Drosophila* species with a history of more intense sexual selection are better adapted to survive environmental stress (e.g., heat waves). However, our data also suggests that the ability of long-sperm species to evolve resilience to heat stress may be constrained by a trade-off with increased susceptibility to heat-induced infertility. Hence, this system highlights that much more work is required to understand how variation in mating systems and the intensity of sexual selection relate to the vulnerability of species in the face of rapid environmental change.

## Acknowledgments/Funding

AMR was funded by an ACCE NERC studentship (project 2750819); SW was funded by the BBSRC (BB/V015249/1 and UKRI2969); RAG was supported by the NSF (DEB-2011045); SP was funded by a generous gift from M. and J. Weeden to Syracuse University.

## References

1. Rowe, L. and H.D. Rundle, The Alignment of Natural and Sexual Selection. Annual Review of Ecology, Evolution, and Systematics 52, 499–517 (2021).

2. Hill, G.E., Condition-dependent traits as signals of the functionality of vital cellular processes. Ecology Letters 14, 625–634 (2011).

3. Warren, I.A., et al., A general mechanism for conditional expression of exaggerated sexually-selected traits. BioEssays 35, 889–899 (2013).

4. Lüpold, S., et al., How sexual selection can drive the evolution of costly sperm ornamentation. Nature 533, 535–538 (2016).

5. Pitnick, S., T.A. Markow, and G.S. Spicer, Delayed male maturity is a cost of producing large sperm in Drosophila. Proceedings of the National Academy of Sciences 92, 10614–10618 (1995).

6. Syed, Z.A., et al., Genomics of a sexually selected sperm ornament and female preference in Drosophila. Nature ecology & evolution 9, 336–348 (2025).

7. Godwin, J.L., et al., Experimental evolution reveals that sperm competition intensity selects for longer, more costly sperm. Evolution Letters 1, 102–113 (2017).

8. Parratt, S.R., et al., Temperatures that sterilize males better match global species distributions than lethal temperatures. Nature Climate Change 11, 481–484 (2021).

9. Sales, K., R. Vasudeva, and M.J. Gage, Fertility and mortality impacts of thermal stress from experimental heatwaves on different life stages and their recovery in a model insect. Royal Society Open Science 8, 201717 (2021).

10. Gustavo Luis, V., et al., Impact of heat waves on semen quality: A retrospective study in Argentina between 2005 and 2023. Science of The Total Environment 943, 173813 (2024).

11. Jørgensen, L.B., R.M. Robertson, and J. Overgaard, Neural dysfunction correlates with heat coma and CTmax in Drosophila but does not set the boundaries for heat stress survival. Journal of Experimental Biology 223, (2020).

12. Matsumura, T., et al., Stress-induced organismal death is genetically regulated by the mTOR–Zeste–Phae1 axis. Proceedings of the National Academy of Sciences 122, e2427014122 (2025).

13. Parrett, J.M. and R.J. Knell, The effect of sexual selection on adaptation and extinction under increasing temperatures. Proceedings of the Royal Society B: Biological Sciences 285, 20180303 (2018).

14. Taylor, M.L., et al., Temperature can shape a cline in polyandry, but only genetic variation can sustain it over time. Behavioral Ecology 27, 462–469 (2015).

15. Meena, A., et al., Life stage-specific effects of heat stress on spermatogenesis and oogenesis in Drosophila melanogaster. Journal of Thermal Biology 125, 104001 (2024).

16. Pitnick, S., Investment in testes and the cost of making long sperm in Drosophila. The American Naturalist 148, 57–80 (1996).

17. Kurhanewicz, N.A., et al., Elevated temperatures cause transposon-associated DNA damage in C. elegans spermatocytes. Current Biology 30, 5007–5017 (2020).

18. Fitzpatrick, J.L., A.F. Kahrl, and R.R. Snook, SpermTree, a species-level database of sperm morphology spanning the animal tree of life. Scientific Data 9, 30 (2022).

19. Team, R.C., R: A language and environment for statistical computing. R Foundation for Statistical Computing. https://www.R-project.org/, (2024).

20. Kim, B.Y., et al., Single-fly genome assemblies fill major phylogenomic gaps across the Drosophilidae Tree of Life. PLoS Biology 22, 3002697 (2024).

21. Revell, L.J., phytools 2.0: an updated R ecosystem for phylogenetic comparative methods (and other things). PeerJ 12,16505 (2024).

22. Tung Ho, L.s. and C. Ané, A linear-time algorithm for Gaussian and non-Gaussian trait evolution models. Systematic biology 63, 397–408 (2014).

23. Martins, E. and T. Hansen, Phylogenies and the comparative method: a general approach to incorporating phylogenetic information into the analysis of interspecific data (vol 149, pg 646, 1998). American Naturalist 153, 448– 448 (1997).

24. Pagel, M., Inferring the historical patterns of biological evolution. Nature 401, 877–884 (1999).

25. Symonds, M.R. and S.P. Blomberg, A primer on phylogenetic generalised least squares. Modern phylogenetic comparative methods and their application in evolutionary biology: concepts and practice 105–130 (2014).

26. Shipley, B., Confirmatory path analysis in a generalized multilevel context. Ecology 90, 363–368 (2009).

27. Van der Bijl, W., phylopath: Easy phylogenetic path analysis in R. PeerJ, 6, 4718 (2018).

28. Gonzalez-Voyer, A. and A. von Hardenberg, An introduction to phylogenetic path analysis. Modern phylogenetic comparative methods and their application in evolutionary biology: Concepts and practice 201–229 (2014).

